# Murine Norovirus infection results in anti-inflammatory response downstream of amino acids depletion in macrophages

**DOI:** 10.1101/2021.04.22.441057

**Authors:** Michèle Brocard, Jia Lu, Belinda Hall, Khushboo Borah, Carla Moller-Levet, Frederic Sorgeloos, Dany J.V. Beste, Ian G. Goodfellow, Nicolas Locker

## Abstract

Murine norovirus (MNV) infection results in a late translation shut-off, that is proposed to contribute to the attenuated and delayed innate immune response observed both *in vitro* and *in vivo.* Recently, we further demonstrated the activation of the eIF2α kinase GCN2 during MNV infection, which has been previously linked to immunomodulation and resistance to inflammatory signalling during metabolic stress. While viral infection is usually associated with activation of dsRNA binding pattern recognition receptor PKR, we hypothesised that the establishment of a metabolic stress in infected cells is a proviral event, exploited by MNV to promote replication through weakening the activation of the innate immune response. In this study, we used multi-omics approaches to characterise cellular responses during MNV replication. We demonstrate the activation of pathways related to the integrated stress response, a known driver of anti-inflammatory phenotypes in macrophages. In particular, MNV infection causes an amino acid imbalance that is associated with GCN2 and ATF2 signalling. Importantly, this reprogramming lacks the features of a typical innate immune response, with the ATF/CHOP target GDF15 contributing to the lack of antiviral responses. We propose that MNV-induced metabolic stress supports the establishment of host tolerance to viral replication and propagation.

**Importance:** During viral infection, host defences are typically characterised by the secretion of pro-inflammatory autocrine and paracrine cytokines, potentiation of the IFN response and induction of the anti-viral response via activation of JAK and Stat signalling. To avoid these and propagate viruses have evolved strategies to evade or counteract host sensing. In this study, we demonstrate that murine norovirus controls the antiviral response by activating a metabolic stress response that activates the amino acid response and impairs inflammatory signalling. This highlights novel tools in the viral countermeasures tool-kit, and demonstrates the importance of the currently poorly understood metabolic reprogramming occurring during viral infections.

## Introduction

The accumulation of viral double-stranded RNA (dsRNA) replication intermediates or proteins during infection imposes a major stress on the host cell. In response to this stress, infected cells induce several defence mechanisms that promote cell survival and initiate an antiviral program (1–3). This first line of defence culminates in a global inhibition of protein synthesis while allowing specific translational control instrumental in promoting the NF-κB and type I IFN-dependent antiviral innate immune response induced downstream of virus recognition by pattern recognition receptors (PAMPs (4–8)). It involves finely choregraphed events such as inhibition of the global translation via phosphorylation of eIF2α (P-eIF2α) and the assembly of stress granules to sequester the bulk of cytoplasmic mRNAs (9) leading to the specific translation of components of the integrated stress response (ISR) genetic program downstream of P-eIF2α (10, 11) and to the specific recruitment into actively translating polysomes of NF-κB and IRF3/7 target genes transcripts such as *TNF*, *Il- 6*, *IfnB1* (12).

The *Caliciviridae* family comprises small non-enveloped positive-strand RNA viruses of medical and veterinary importance (13). Among these, Human Norovirus (HuNoV) is a major cause of acute gastroenteritis outbreaks worldwide, responsible for more than 200,000 deaths per year, and has a socioeconomic impact estimated at more than $60 billion/year (14, 15). Both HuNoV and murine norovirus (MNV) belong to the Norovirus genus and share many characteristics, with MNV benefiting from reverse genetics systems, small animal model, and easy propagation in cell culture (16). Therefore, MNV provides a valuable model for understanding the life cycle of caliciviruses.

MNV replication is sensed by melanoma differentiation-associated protein 5 (MDA5, (17)), a member of the RIG-I-like receptors family (RLRs), leading to a robust IFN response (18), which however fails to contain MNV replication and propagation. Moreover, while MNV replication is susceptible to IFN pre-treatment or activation of the Toll-like Receptor (TLR) cascades, these antiviral responses become ineffective after the early phase of infection, suggesting escape from antiviral responses (19, 20). Furthermore, our previous studies have shown that MNV infection regulates translation in several ways: by controlling the activity of multiple eIFs by inducing eIF4E phosphorylation via the MAPK pathway, and by cleavage of PABP and eIF4G by the viral protease or cellular caspases (19, 21). We and others also recently showed that while MNV impairs global translation, this shutoff is uncoupled from the activation of the eIF2α-dependent stress responses (19, 21–23).

In response to viruses, the ISR is classically linked to the activation of the eIF2α kinases PKR by viral dsRNA replication intermediates, or PERK due to the accumulation viral proteins in the ER (10, 24, 25). However, our previous study identified the kinase GCN2 as responsible for MNV-induced phosphorylation of eIF2α. GCN2 is activated by numerous stresses, including amino acid starvation and ribosomal stresses such as translation elongation defects and ribosomes collisions, triggering the ISR downstream of translation inhibition (26–28). Interestingly, several lines of evidence suggest a strong link between metabolic homeostasis, immunotolerance and immunosuppression correlating with GCN2 activity (29–31). Sensing of metabolic stress also initiates the Immediate Early Response (IER), leading to activation of the transcription factor ATF2 via the MAPK JNK (32). The resulting response to both the IER and ISR, also called the amino acid response (AAR) revolves around the transcriptional activity of the transcription factors ATF4, ATF3 and CHOP leading to a homeostatic response to the identified stress and suppression of NF-κB - dependent inflammation or to apoptosis via the intrinsic apoptosis pathway if this stress cannot be resolved (10). Noticeably, ATF3 had been shown to negatively regulate the transcription of pro-inflammatory genes such as *Il-6* and *Il12p40* (30, 33). This, together with the links between GCN2 activity and immunosuppression, and its antiviral roles during Sindbis virus or HIV infection (6, 29–31, 34), puts GCN2 at the nexus between MNV infection, stress responses and innate immunity.

Herein, we hypothesise that the impaired antiviral response during MNV infection may in part originate from a metabolic stress activating GCN2-mediated ISR and IER, leading to the suppression of the NF-κB -dependent inflammation downstream of MDA5 activation. To test this, we identified an imbalance in amino acid metabolism in MNV-infected cells and characterised the cellular response using genome-wide analysis of the transcriptome, translatome and proteome in MNV- infected macrophages cell lines. We demonstrated the activation of an MDA5- dependent pathway lacking features of a typical innate immune response and correlating with the activation of ISR / IER genetic program during MNV infection. We further show the importance of ATF3 upregulation, and the ATF/CHOP target Gdf15, a tolerogenic factor and non-steroidal anti-inflammatory drugs activated gene (NAG- 1) (35), in controlling paracrine inflammatory targets such as Ptgs2 (Cox-2) or Mx1. Altogether, our results highlight a previously undescribed mechanism of control of the anti-viral response in macrophages via activation of a metabolic stress response associated with MNV-induced tolerance.

## Results

### MNV induces an amino acid imbalance in infected cells

We previously identified a role for GCN2 in inducing eIF2a phosphorylation during MNV infection (23). Given the link between GCN2 activation and the response to amino acid starvation, we assessed amino acids levels in MNV-infected cells by first measuring the total content in free amino acids using a colorimetric enzymatic assay in mock- or MNV-infected RAW264.7 murine macrophage cells at 10h p.i. using cells grown in Glycine-Cysteine-Methionine (GCM) -depleted medium as control. The results showed no significant differences in the total concentration of free amino acids between all conditions suggesting the absence of global amino acid depletion during infection (Fig. 1A). Next, we quantified the pool sizes of individual amino acids using Gas Chromatography-Mass Spectrometry (GC-MS). This analysis revealed significant differences in the pool sizes of free amino acids in MNV-infected cells, interestingly with an overall similar pattern similar to that observed for cells grown in GCM depleted medium (Fig. 1B). Specifically, GCM-depletion increased the glutamine, glutamate, serine and leucine and reduced the methionine, cysteine and phenyalanine pool size, which was mirrored in MNV-infected cells suggesting that viral infection induced a similar metabolic stress response as amino acid starvation. Additionally, virally infected cells had increased pool sizes of free valine, isoleucine, proline and aspartate. Although we cannot make any conclusions about specific changes in metabolic flux from these results, this data confirms that MNV replication induces a metabolic stress response which disrupts amino acid homeostasis. Beside GCN2 activation, the amino acid response is characterised by the activation of the MAPK pathway, specifically its JNK2 arm, leading to the activation of the AP1 transcription factor ATF2 (32, 36). Our previous analysis in MNV-infected RAW264.7 cells showed the activation of Erk1/2 and p38 activation, as part of a classical anti- viral response (21). Similar MAPK antibody arrays were used to examine the phosphorylation of JNK isoform at 2 and 12h p.i. in mock- and MNV1-infected RAW264.7 cells. These data, summarized in Fig. 1C, show at both 2 and 12h p.i. an increase in phosphorylated JNK2, and to a lesser extend of JNK1, in MNV-infected cells. These results further support the sensing of a metabolic imbalance and activation of the AAR, encompassing the observed activation of GCN2 on one hand and the activation of JNK pathway on the other. Next, we used immunoblotting to examine the phosphorylation status of ATF2 at Thr71 in MNV-infected cells (Fig. 1D). We then confirmed by qRT-PCR the upregulation of the ISR and IER response genes *ATF3* and *Ddit3* in MNV-infected cells (Fig. 1E and F), which can be partially reverted by the GCN2 inhibitor A92, at a concentration known to inhibit the GCN2- dependent phosphorylation of eIF2α in MNV-infected cells (23). Altogether, these results suggest the activation of AAR signalling downstream of an amino acid imbalance, culminating in the expression of its associated genetic program during MNV infection.

**Fig. 1:**
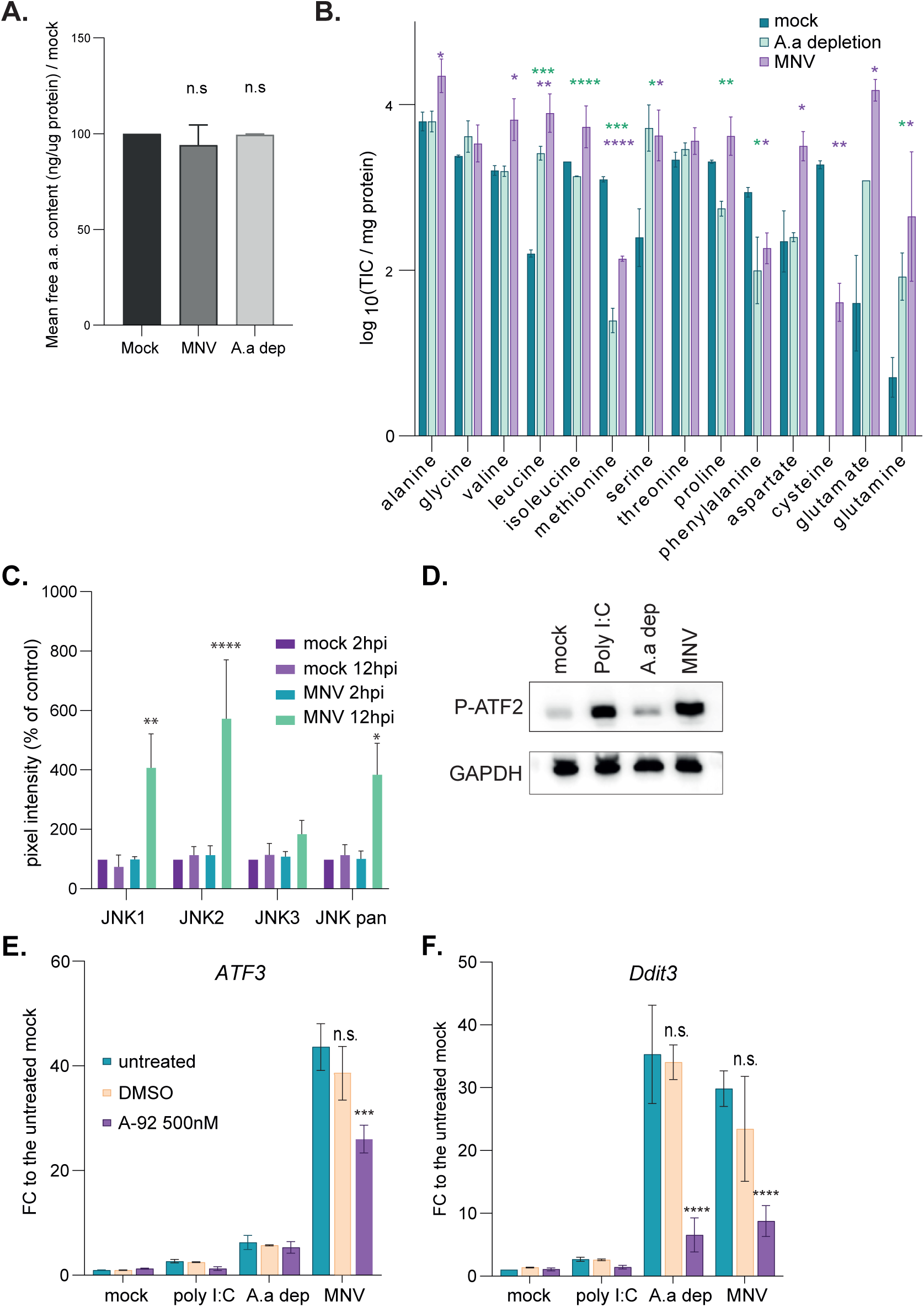
MNV infection is associated with an amino acid depletion phenotype. (A and B) Quantification of free amino acids in RAW264.7 cells mock- (dark green), MNV-infected (magenta) for 10h or incubated in GCM depleted medium (light green). There were no significant differences between the total concentration of free amino acids, quantified using a colorimetric assay. By contrast GC-MS (B) measured significant differences in the pool sizes of individual amino acids (TIC, Total Ion Current). Values are mean ± s.e.m (n=3), submitted to *t-test* pairwise comparison, significance **** *P* < 0.0001, *** *P* < 0.001, ** *P* <0.01, * *P* < 0.1. The mass-to-charge ratios for analysed amino acids are alanine 260, glycine 246, valine 288, norvaline 288, leucine 274, isoleucine 274, methionine 320, serine 390, threonine 404, proline 258, phenylalanine 336, aspartate 418, cysteine 406, glutamate 432, ornithine 417, asparagine 417, lysine 431, glutamine 431, arginine 442, histidine 440, tyrosine 466, tryptophan 375. (C) Stress related JNK1/2 are activated during MNV replication in RAW264.7 cells. Bar plot (n=3) of the mean ±S.D. of the results of phospho-antibody array assay (n=3) at 2h p.i. (light) and 12h p.i. (dark) in mock- and cells infected with MNV at MOI 10 (respectively green and purple bars). Statistical analysis results shown above the plots, * *P*<0.1, ** *P*<0.01, **** *P*<0.0001. (D) Western blot analysis of the activation of ATF2 in mock cells, cells treated with 20µg/ml of poly I:C, amino acids starved cells and MNV-infected cells at 10h p.i. (MOI 10). (E and F) GCN2 is partially responsible for the genetic reprogramming in MNV-infected cells. Bar plots (n=4) of the mean ± s.e.m of transcripts upregulation analysis by qRT-PCR for (E) *ATF3* mRNA and (F) *CHOP* mRNA. Experiment performed on total transcripts from RAW264.7 cells treated as in for 10h, with or without 500nM of GCN2 inhibitor A92. Purified RNAs were reverse transcribed using polydT primer and qPCR performed using exon-junction spanning pair of PCR primer. Statistical significance between untreated and treated samples calculated using Anova 2way multiple comparison tool in GraphPad shown above the bars, **** *P* < 0.0001, *** *P* < 0.001, n.s. not significant.

Genetic reprogramming analysis reveals an AAR response in MNV-infected cells. The homeostatic response AAR is responsible for starvation-induced survival and suppresses any further inflammation via the ATF4 and ATF2 response gene *ATF3* (37, 38), partly by inhibiting the upregulation of NF-κB target genes *Il-6* and *Il12p40* (30, 33). To understand the global impact of this amino acid imbalance on the host anti-viral response to MNV infection, we performed a genome-wide analysis of the transcriptome and translatome. RNA-Seq analysis of the cytoplasmic polyA-tailed transcripts (total fraction) and polyA-tailed transcripts associated with polysomes (polysomal fraction) was carried out at 6 and 10h p.i. corresponding to time-points prior to and during the MNV-induced phosphorylation of eIF2α by GCN2 (23), in RAW264.7 infected at MOI 10 with matching controls using UV-inactivated virus, MNV(UVi) (Fig. 2A). Transcripts associated with polysomes were isolated after ultracentrifugation through sucrose gradients and pooling of the fractions 5 to 10 (Fig. S1A to S1C). Three independent replicates were used, with similar viral replication efficacy across replicates as analysed by TCID50 (Fig. 2A insert). The extracted RNA samples were evaluated for concentration and quality with a Bioanalyzer before analysis by RNA-Seq. All sample reads were aligned to murine genome assembly GRCm38.p5 using Hisat2 and MNV1-CW1 genomic RNA (DQ285629) using Bowtie2 (Fig S1D and E). This revealed that 98.2%±0.2% and 97.4%±0.06% of reads aligned to the host genome in total and polysomal fractions respectively at 6h p.i., and this decreased to 69.9%±1.7% and 71.4%±1.4% at 10h p.i., reflecting increased viral RNA synthesis and translation (Fig S1E). Bi-orthogonal plotting of the averages of normalised reads (log_2_CPM) of the three replicates for the different conditions showed a striking relationship between polysomal and total cytoplasmic fractions using nonparametric Spearman correlation test (ρ>0.98, pval<0.0001 across all the conditions), supporting a global transcript recruitment into the polysomes as a function of its cytoplasmic availability even in MNV-infected samples at 10h p.i. (Fig. 2B to 2E). Further filtration of the datasets by gene feature annotation led to an enrichment for the protein coding transcripts of 85.9% (Fig. S1F). Correlation analysis showed strong similitude (Pearson Correlation Coefficient r>0.98) with clear hierarchical clustering following the parameters: infection, time, fractions and no predominant batch effect (Fig. S1G). As differential expression analysis requires compositional normalisation and considering the vast amount of MNV RNA in the 10h p.i. samples, we created sub-datasets normalised by library size omitting the MNV RNAs (Host dataset). 3D principal-component analysis (PCA) of the complete dataset (MNV + host genes) showed an expected clustering of the MNV-infected samples (PC1) over other parameters. Similar analysis on the host dataset while showing a more complex relationship between samples, nevertheless highlighted a clear clustering of the MNV samples at 10h p.i. following mainly the fraction parameter (PC1) and a resultant of infection, time and replicates parameters (PC2 and PC3) (Fig. S1H and I).

**Fig.2:**
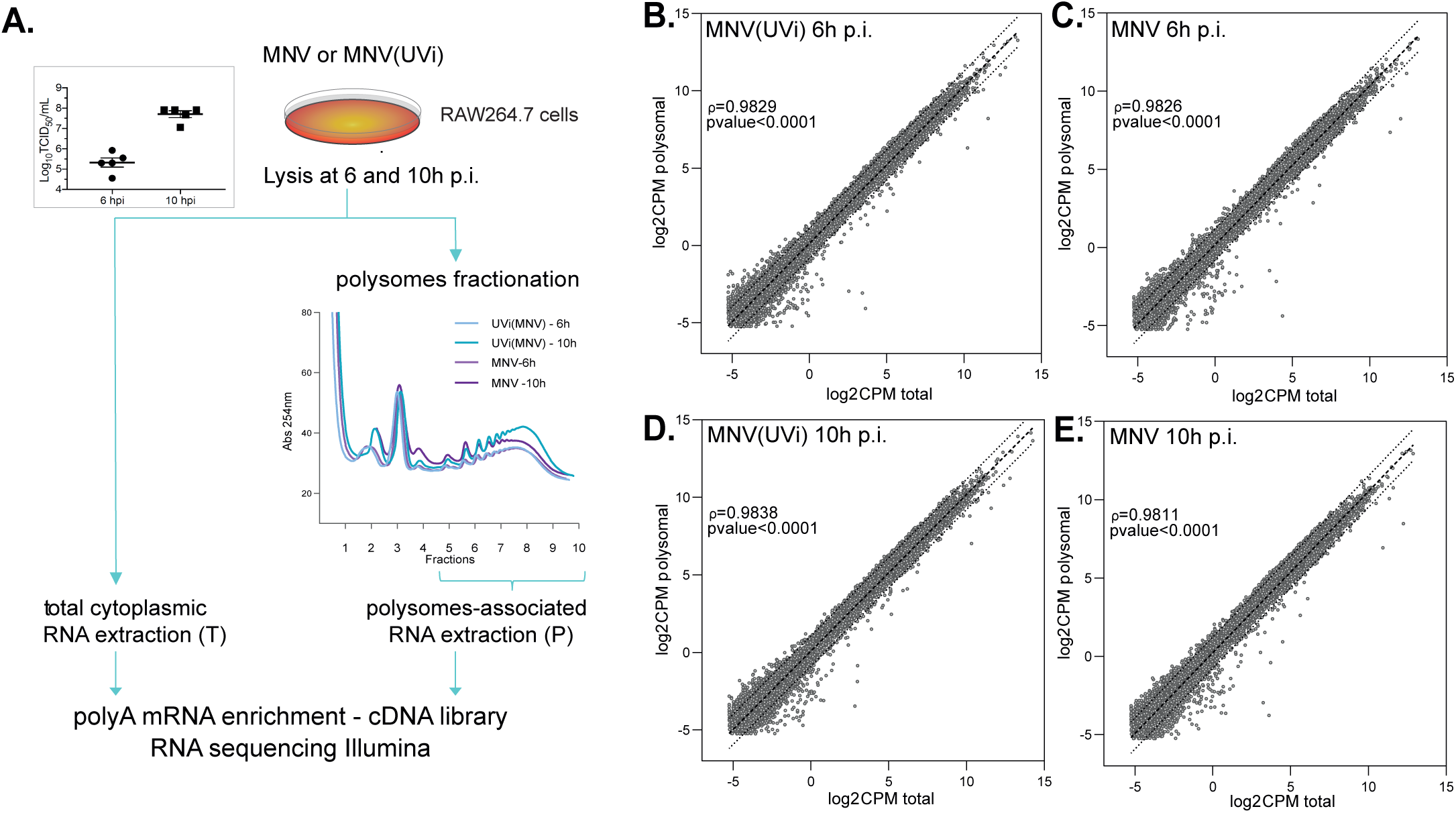
Multi-variate omics analysis of MNV-infected cells by RNA sequencing. Comprehensive genome-wide RNA sequencing analysis comparing the changes of the landscape of cytoplasmic transcripts available for translation (Total fraction) to the corresponding changes of the landscape of transcripts recruited into polysomes (Polysomal fraction), their kinetic variation from 6 to 10h p.i., in murine macrophages cell line RAW264.7 infected with the replicative MNV (MOI 10) or the non-replicative UV-inactivated MNV (MNV(UVi)). (A) Diagram of the experimental design for the translatomic analysis experimental design, from infection to RNA sequencing, also showing in insert the similar efficiencies of MNV infection across samples and replicates addressed by measure of the viral titre by TCID50 (logarithmic scale) at 6 and 10h p.i. (B. to E.) Scatter plots of the RNA sequencing results plotted as the average log_2_CPM in Polysomal (*y* axis) vs Total fraction (*x* axis) for each condition, MNV(UVi) (B and D) or MNV(C and E) at 6 (B and C) or 10h p.i. (D and E). The log_2_CPM of the host dataset genes were obtained by library size normalisation of the raw results and filtering out of the low reads genes (CPM<8). Correlation analysis using the ‘Simple linear regression’ function from GraphPad showing the strong relationship (Spearman correlation coefficient *ρ*>0.98, pval<0.0001) between the Total and Polysomal fractions. Dotted lines, linear regression and 95% confidence intervals.

Initial differential expression analysis in total and polysomal fractions between MNV and MNV(UVi) conditions at 6 and 10h p.i. highlighted a strong overlap of behaviour for the majority of genes as suggested by their projection membership into the 95% interval of confidence of significance in both conditions (Fig. S2A and S2B). This overlap likely reflects the physiological changes that occur to the cells during the experiment rather than a direct response to infection per se. The heatmap of differentially expressed genes between 6 and 10h p.i. in MNV-infected samples highlights a lack of clear difference between MNV and MNV(UVi)-infected samples with a confused clustering following infection, time and fraction parameters (Fig. S2C), suggesting weakening of the statistical analysis due to the introduction of too many variability parameters between isogenic samples and increasing the noisiness of the results. We therefore performed the more robust comparison of the samples prepared temporally in a parallel manner: MNV *vs* MNV(UVi) at 6 and 10h p.i. While we did not detect any significant differentially expressed genes at 6h p.i. between MNV- and MNV(UVi)-infected cells (Fig. S2D), we identified 265 differentially expressed genes in the total fraction and 229 differentially expressed genes in the polysomal fraction (Fig. 3A) at 10h p.i., all of which displayed a p-value inferior to 0.0025 for BHpval<0.1 (Fig. 3B and C). Validation of those results by RT-qPCR showed a significant correlation (Spearman ρ= 0.9564, pval<0.0001) for the assayed genes (Fig. S2E). While 67 genes showed a significant differential expression only in the total fraction and 31 genes only in the polysomal fraction, the majority fall into a 95% prediction band of co-regulation which seems to confirm the recruitment of transcripts to the polysomes fraction according to their abundance. A hierarchical clustered heatmap of the Log_2_CPM of all the significant differentially expressed genes across the replicates confirmed the strong influence of the infection parameter as the source for differences of expression (Fig. 3D). Moreover, translational efficiency analysis performed with the R package Anota2seq (39) highlighted a lack of general translational control in MNV-infected cells at 10h p.i. (Fig. 3E), leaving only the partial translation shutoff previously described.

**Fig.3:**
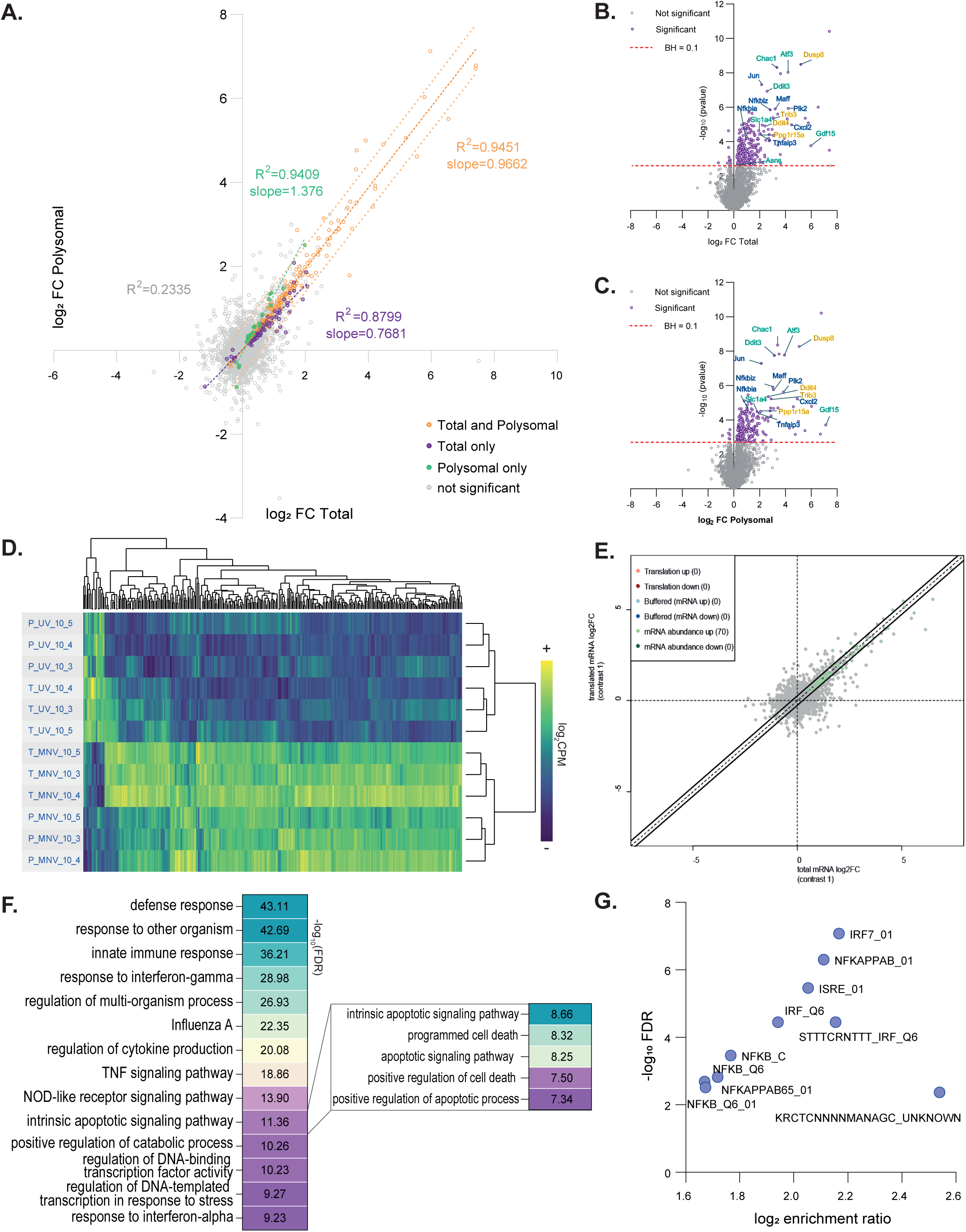
MNV infection induces an attenuated innate immune response as seen by RNA sequencing of both total and polysomal fractions in RAW264.7 cells at 10h p.i. Differential analysis performed on RNA sequencing data showed a genetic reprogramming induced by MNV replication in RAW264.7 cells lacking both NF-κB and Jak/Stat response. (A) Scatter plot of the results of the differential expression analysis comparing MNV-infected cells vs MNV(UVi)-infected cells at 10h p.i. in the Total (*x* axis) vs the Polysomal fraction (*y* axis) showing mainly a subset of MNV- specific upregulated genes with similar behaviour in both fractions. The differential expression analysis was performed on all samples at 10h p.i. using the multivariate tool GLMEdgeR and gene level changes were considered significant when associated with a BHpval <0.1. Results expressed as log_2_ (FC MNV vs MNV(UVi)) of the complete gene dataset were used for the biorthogonal projection and subdivided into four groups: genes significant in both Total and Polysomal fractions (orange dots), genes significant in Total fraction only (purple dots), genes significant in Polysomal fraction only (green dots), and genes displaying no changes (grey dots). Correlation analysis using the Pearson method showed high degree of correlation for the three significant groups as indicated on the plot. Linear regression (full orange line) and prediction bands at 95% confidence of significance for the “significant in both fractions” genes (dotted orange lines) seemed to encompass most of the genes significant in one fraction only reflecting the stringency of the significance cut-off and the likeliness of changes in both fractions for all the significant genes. (B and C) Volcano plots of the log_2_FC (*x* axis) vs the corresponding -log_10_(pval) (*y* axis) for each gene of the Total (B) or the Polysomal dataset (C) from the differential expression analysis MNV vs MNV(UVi)-infected cells at 10h p.i showing the high degree of significance (Total pval<0.0025, Polysomal pval<0.002) resulting from the BHpval<0.1 thresholding. Threshold of significance BHpval<0.1 (red line), significant genes (purple dots), non-significant genes (grey dot). Classical target genes downstream of the ISR via ATF4 (yellow), downstream the IER via ATF2 (blue) and via both ATF4 and ATF2 in response to amino acid starvation (green) (D) Heatmap of comparison of the significant differentially expressed genes in MNV-infected cells displaying their log_2_CPM values in each sample showing a strong clustering following the parameter infection and fraction but not set of biological replicates. Log_2_CPM values of significant genes for all samples were centred and scaled, subjected to correlation analysis using the Pearson method and complete linkage hierarchical clustering. (E) Scatter plot of the translational control analysis results showing an absence of translational regulation in MNV-infected cells, Translated mRNA (*y* axis) vs Total mRNA (*x* axis). (F) Clustered results of the gene ontology analysis showing enrichment of MNV-induced upregulated genes in anti-viral innate immune response terms and intrinsic apoptotic pathway. Insert showing the different terms associated within the “Intrinsic apoptotic pathway” cluster. The gene annotation analysis was performed on Biological Process and KEGG-pathways terms using the Cluego plug-in on the Cytoscape platform for all genes significantly regulated at 10h p.i. (Total and Polysomal dataset) and ordered by decreasing corresponding -log_10_(FDR). (G) Scatter plot of the transcriptional network analysis results for the MNV-induced upregulated genes at 10h p.i. showing enrichment for the IRF, NF-κB and IFN-Response genes pathway. Enrichments were calculated on the transcription factor target database MSigDB using the over-representation analysis method and plot created by biorthogonal projection of the log_2_ enrichment ratio (*x* axis) vs the -log_10_(FDR) (*y* axis).

Interestingly, most of the identified differentially expressed genes are upregulated (281 out of 296), which is in stark contrast with previous published transcriptomic data of MNV-infected cells (40, 41) and could reflect the bias introduced by the comparison of temporally unmatched conditions in previous studies (*i.e.* MNV 12h p.i. vs mock or MNV 0h p.i.) and low level of infection. Direct comparison showed 31 genes upregulated in all 3 studies in the total fraction, which are quite likely to be the core signature of MNV infection and illustrate the absence of downregulated genes, likely to be linked to physiological changes over time in the previous studies (Fig. S3A). Functional annotation analysis and group clustering using Cytoscape (42) (Fig. 3F and S3B) showed enrichment of the upregulated genes for an apparent classical anti-viral response with “defence response to organism” (-log_10_(FDR)=43.11), ”response to other organisms” (-log_10_(FDR)=42.69), “Innate immune response” (- log_10_(FDR)=36.21), “response to cytokine” (-log_10_(FDR)=34.87), “response to interferon gamma” (-log_10_(FDR)=28.98), “Influenza A” (-log_10_(FDR)=22.35) or “TNF signalling” (-log_10_(FDR)=18.86). These results support the 2 previous transcriptomic analysis showing the activation of MAPK kinase and pathways downstream of MDA5 activation during MNV replication and confirmed by transcription target factor analysis and enrichment for the Interferon Regulatory Factors (IRF) and Interferon Stimulated Response Element (ISRE) target genes (Fig. 3G). In addition, MNV- induced differentially expressed genes are enriched for “positive regulation of catabolic process” (-log_10_(FDR)=10.26), which matches the observed MNV induction of autophagy (43). Genes upregulated in MNV-infected cells also showed a strong enrichment for “intrinsic apoptotic signalling pathway” (-log_10_(FDR)=11.36), encompassing different cell death-related GO terms but only mentioning the intrinsic aspect (-log_10_(FDR)=8.67, Fig. 3F insert), fitting the known activation of the intrinsic apoptotic pathway in MNV-infected cells (44). The 15 downregulated differentially expressed genes did not show any enrichment using the same setting, however a more relaxed metascape-driven analysis highlighted an enrichment in “positive regulation of cell cycle” (*i.e. E2F2* and *E2F7* -log_10_(pval)=4.21), also fitting a previous publication on the effect of MNV on the mitotic activity of the infected cells, especially the G1 to S phase transition (45).

Interestingly, the genes enriched in “NOD-like receptor pathway” are the IER target genes downstream of ATF2 activation (*Tnf, NFκBia, Tnfaip3*, *Cxcl2* (46)). Moreover, those enriched in the “intrinsic apoptotic signalling pathway” (-log_10_(FDR)=11.36), and “regulation of DNA-templated transcription in response to stress” (- log_10_(FDR)=9.27) are known ISR-induced genes such as *ATF4*, *ATF3*, *Ddit3* (CHOP), *Chac1*, *Ppp1r15a* (GADD34), *Trib3*, *Bcl2l11* (Bim), *Dusp2* and *Dusp8* and belong to the ATF4-CHOP apoptotic pathway described downstream of the ISR (10, 11, 47). Importantly, the upregulation of *Ddit3* expression (Total_log_2_FC= 2.57) in absence of *Hspa5* upregulation (Total_log_2_FC= 0.15) is classically linked to nutrient starvation in the absence of Unfolding Protein Response stimulation (48), which also points to the identified metabolic stress. Furthermore, the upregulation of *Ddit3* transcripts confirmed the activation of both the ISR and ATF2 pathways concomitantly as both are necessary for its transcriptional activation (49). These results confirm the induction of the AAR genetic program in MNV-infected cells downstream of both the ISR and the IER.

In contrast, a comparison with the previously identified LPS-induced genes in RAW264.7 cells (12) showed a strong defect in the cytokines induction required for autocrine/paracrine establishment of an anti-viral environment downstream of NLR and TLR activation (4, 7, 8). Specifically, we observed no upregulation of pro- inflammatory chemokines and cytokines coding genes such as *Il-6, Il-18, Il-1α, Il-1β, Il-23, Ccl5* or *Il-12p40* (7) (Fig. 4A), known NF-κB target genes, and *Csf2* and *Ptgs2* linked to Stat5 transcriptional activity in bystander cells (50). Transcriptional network analysis of these genes further highlights the strong enrichment in NF-κB binding motifs in their promoters (Fig. 4B). Moreover, the expected programmed cell death downstream of NLR and TLR via the extrinsic apoptotic pathway, necroptosis and pyroptosis (51) and depending of NF-κB induction of *Fas or Tnfrsf10b*, also seems to be absent in MNV-infected cells (Fig. 4C). Altogether, these results reflect the known activation of MDA5 in MNV-infected cells but indicate an inappropriately attenuated NF-kB activity downstream of MDA5 further suggesting a suppressive mechanism correlating with the induction of the anti-inflammatory AAR program.

**Fig.4:**
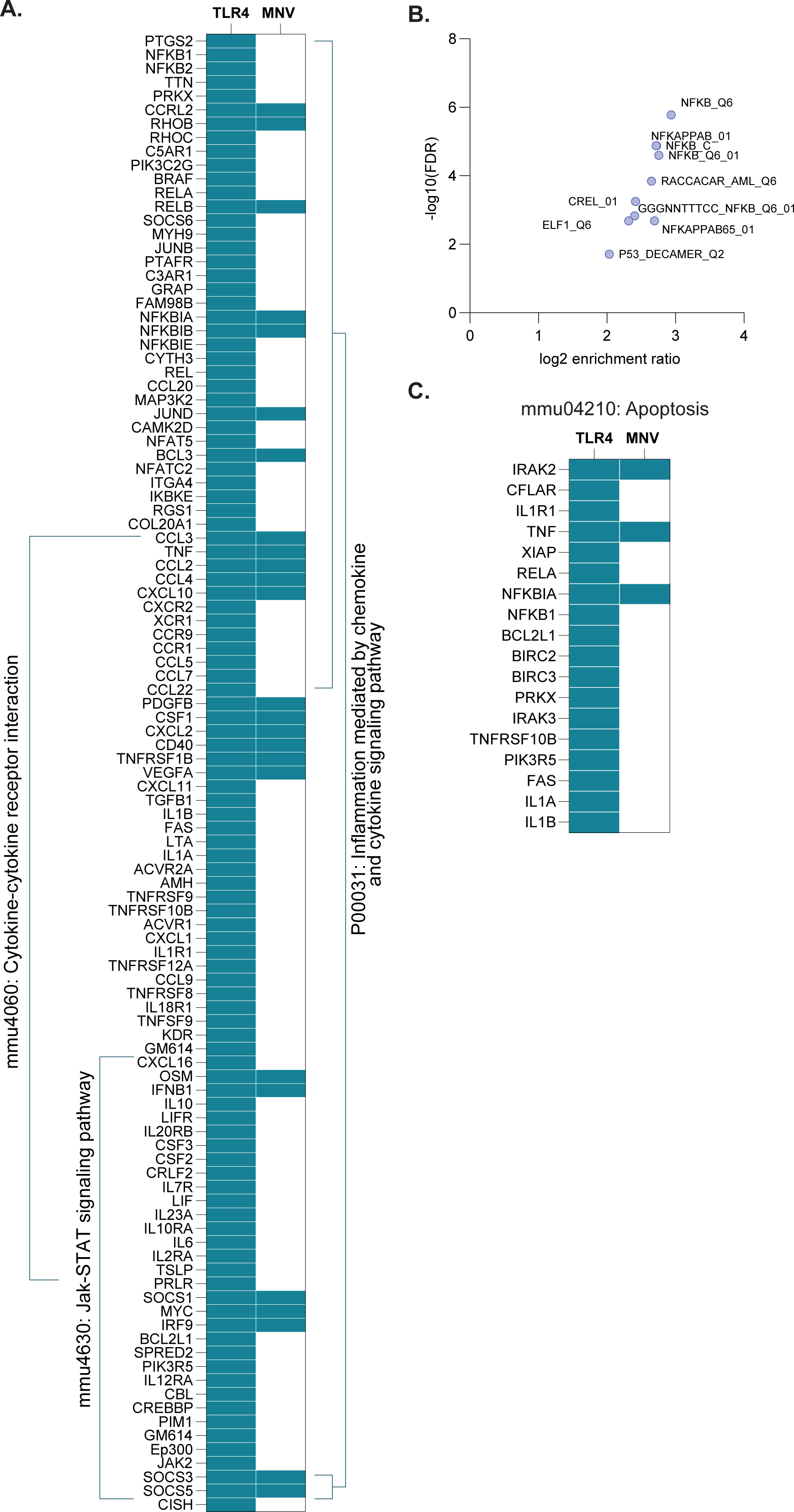
Host response to MNV is lacking crucial pro-inflammatory cytokines, targets of JAK/STAT signalling and inflammation-related cell death factors linked to autocrine and paracrine activation of the innate immune response. (A) Comparison of the differentially expressed genes associated with the LPS activated TLR4 response in RAW264.7 cells with the significantly regulated genes in MNV-infected RAW264.7 cells at 10h p.i. Significantly differentially expressed gene, blue square, absence of differential expression, white square. List of cytokines had been generated from the list of differentially expressed genes (group 1) following LPS activation of RAW264.7 cells from (12) related to the pathway terms “mmu04060_Cytokine-cytokine receptor interaction”, “mmu04630_Jak-STAT signalling pathway”, “P00031_Inflammation mediated by chemokine and cytokine signalling pathway” generated using KEGG_pathways and Panther tools on the Cytoscape platform, gene symbols had been matched to fit the more recent annotation and run against the list of significant differentially expressed genes in MNV-infected cells at 10h p.i. using the open- source JVenn tool. (B) MNV-infected cells are lacking activation of genes related to NF-κB transcriptional network. Scatter plot of the transcriptional network analysis results for the “TLR4 only” genes regulated after TLR4 activation and not regulated in MNV-infected cells. “TLR4 only” list of genes generated from the JVenn comparison as in (A). Enrichments were calculated on the transcription factor target database MSigDB using the over- representation analysis method and plot created by biorthogonal projection of the log_2_ enrichment ration (*x* axis) vs the -log_10_(FDR) (*y* axis). (C) Comparison of the differentially expressed genes associated with the TLR4 response and enriched in “mmu04210_Apoptosis” pathway” KEGG pathway. Significantly differentially expressed gene, blue square, absence of differential expression, white square. List of genes generated as in (A).

### Antiviral and anti-inflammatory pathways are both activated in MNV-infected macrophages

We therefore addressed the kinetic of upregulation of *ATF3*, *TNF*, *Il-6* and *IFNβ1* genes expression in MNV-infected cells compared with poly(I:C) activated cells by qRT-PCR (52). *ATF3* expression is strongly activated between 6 and 8h p.i. prior or concomitantly to the upregulation of *IFNβ1* and outstandingly in the absence of priming *TNF* upregulation which is only seen for the poly(I:C) treated cells (Fig. 5A to D). This confirms that activation of a proinflammatory signalling pathway by cytokines-mediated amplification loops is missing in MNV-infected cells prior to the modulatory ATF3 response. Rather, MNV replication seems to induce ATF3 dependent anti-inflammatory signalling at the same time as the MDA5-mediated cellular innate immune response.

**Fig.5:**
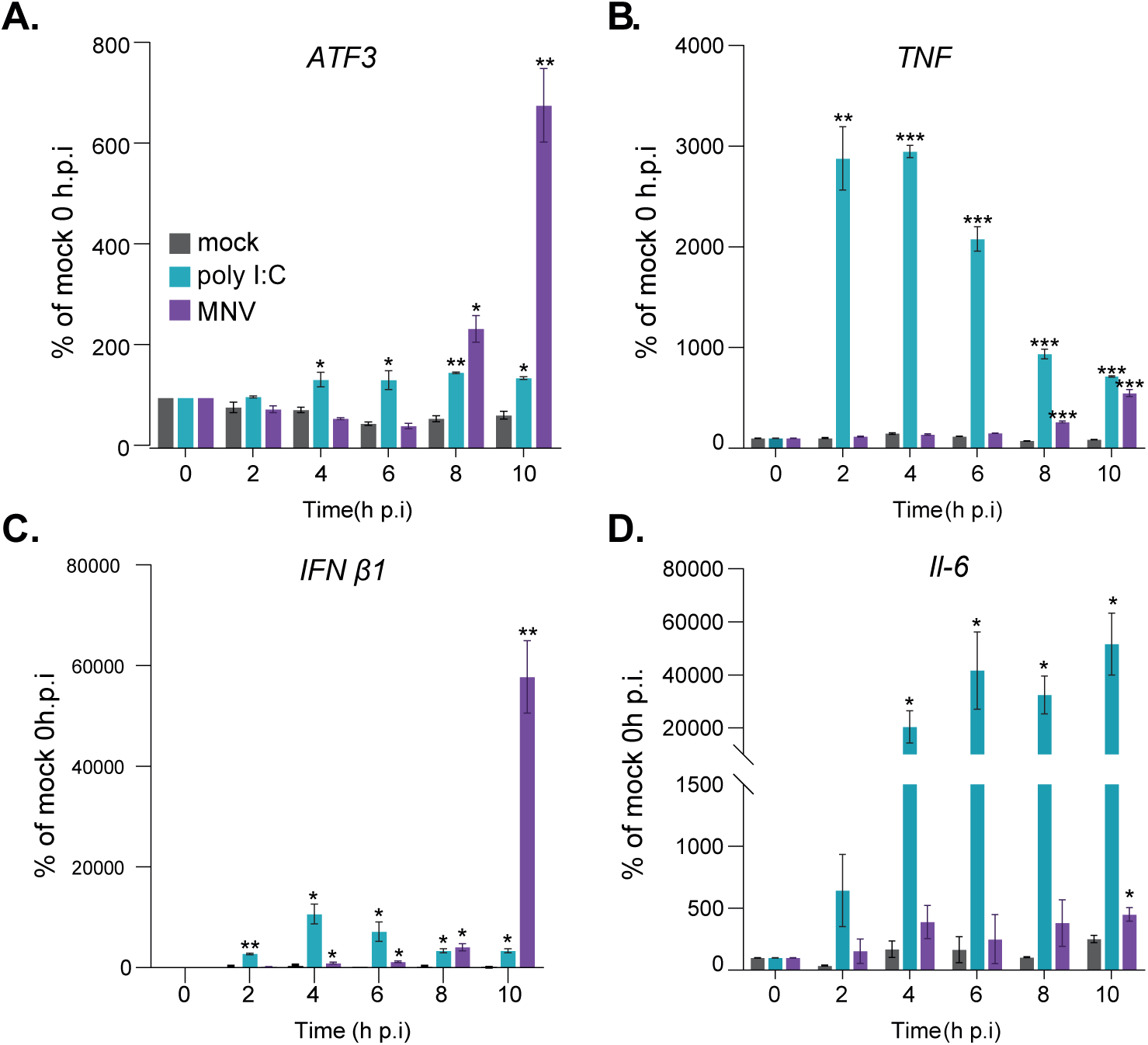
ATF3 response is not secondary to MDA5 response during MNV infection. qRT-PCR analysis of a time-course experiment of RAW 264.7 cells mock (grey), poly I:C treated (cyan) or infected with MNV (purple) at MOI 10 for 10h. Representative results (n=3) as mean ± S.D. showing the kinetic of regulation of (A) *ATF3* mRNA, (B) *TNF* mRNA, (C) *Ifnβ1* mRNA and (D) *Il-6*mRNA. Statistical analysis results shown above the plots, * *P*<0.1, ** *P*<0.01, *** *P*<0.001.

Next, we confirmed the ability of infected cells to actively translate this transcriptional reprogramming. We measured temporal changes in the nascent proteome by biorthogonal noncanonical amino acid tagging (53), followed by mass-spectrometry in cells labelled with the amino acid analogue L-azidohomoalanine (AHA) at 6 and 10h p.i., comparing mock, MNV(UVi) and MNV infected cells (Fig. 6A). This analysis identified 2,661 host and 6 viral proteins at 6h p.i. and 3,396 host and 9 viral proteins at 10h p.i. Pairwise ratio comparison identified 56 proteins differentially translated at 6h p.i. (Fig. S4) and 89 proteins at 10h p.i. when comparing both MNV vs mock and MNV vs MNV(UVi) (Fig. 6B). No specific enrichment was found for the 19 upregulated proteins at 6h p.i. but the 37 downregulated proteins were enriched for “Positive regulation of mitotic nuclear division” (3 genes, -log_10_(FDR)=3.67). Functional annotation analysis of the 10h p.i. results (Fig. 6C) revealed an enrichment of the 52 upregulated genes in “defence response to virus” (- log_10_(FDR)=13.11), with several IFN type I related GO terms, showing the solid transactivation of this response: “regulation of type I Interferon production” - log_10_(FDR)=10.87, “response to Interferon-beta” -log_10_(FDR)=9.15, “response to Interferon-alpha -log_10_(FDR)=6.34), but there was no enrichment in TNF nor NFκB pathways. The 37 downregulated proteins were enriched in “Regulation of cyclin- dependent protein serine/threonine kinase activity” (-log_10_(FDR)=4.6) which could also contribute to the observed effect of MNV on the cell cycle (45) (Fig. 6D). Comparative analysis with the RNA-Seq datasets (Fig. 6E, 83 genes) showed a positive correlation of behaviour for a subset of upregulated genes (Pearson r=0.5943, pval<0.005) belonging to the IFN response such as *Usp18*, *Irgm1* or *Ddx58* (MDA5), confirming the translation of a specific transcriptional programme in MNV-infected cells, albeit reduced as shown previously (23), and an autocrine IFN response. A subset of genes was significantly translationally upregulated but showed no significant differential expression in the RNA-Seq analysis, reflecting differences of stringency between both methods (i.e. *Parp14* and *Dhx58*) or specific changes in protein stability during infection (i.e. *Tmem259*). Downregulated proteins showed no correlation with the RNA-Seq analysis (i.e *Tm9sf1* and *Trappc8*) suggesting a decrease in stability for those proteins. The absence of obvious targets of the ATF4/ATF3 pathway among the proteins identified by SILAC prompted us to use immunoblotting to assess the upregulation of ATF3 and inhibition of TNF expression during infection when compared to poly(I:C) treatment (Fig. 6F). These results support the activation of an immunomodulatory pathway and an attenuated NF-κB response, together, rather than downstream of, in response to MNV infection.

**Fig.6:**
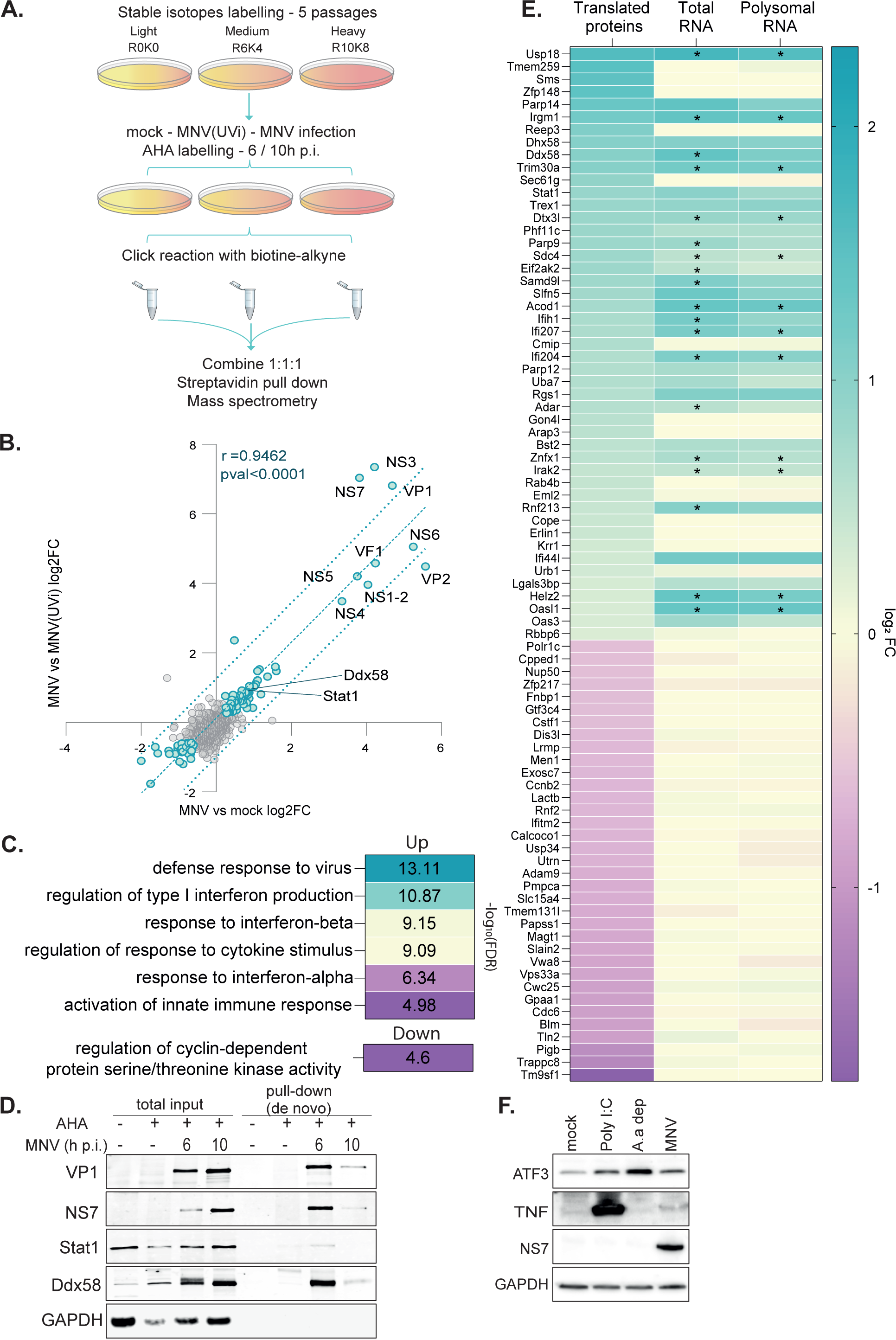
Validation of the results of the translatomic data analysis by mass spectrometry analysis of the nascent proteome in RAW264.7 cells. Genome wide analysis of the nascent proteome by biorthogonal labelling of the nascent protein and mass spectrometry allowing an extensive validation of the active translatome identified by RNA sequencing (A) Diagram of the experimental design of the biorthogonal dual labelling proteomics assay as described in (53) from SILAC labelling to mass spectrometry. (B) Scatter plot of the results of the differential expression analysis in the contrasts MNV- / mock-infected cells (*x* axis) vs MNV- / MNV(UVi)-infected cells (*y* axis) at 10h p.i. showing the enrichment for viral structural and non-structural proteins as well as a subset of MNV-induced differentially expressed host proteins with similar behaviour in both contrasts as shown by the correlation analysis. The differential expression analysis was performed on all samples at 10h p.i. using pairwise ratio and results expressed as log_2_ (FC) of the complete gene dataset were used for the biorthogonal projection, significantly regulated protein (green dots), not significantly regulated protein (grey dots). Correlation analysis using the Pearson method showed high degree of correlation between the two contrasts (*R^2^*>0.89, pval<0.0001). Dotted blue line, linear regression and IC95% of significance. (C) Clustered results of the gene ontology analysis showing enrichment of the upregulated host proteins in anti-viral innate immune response terms and enrichment of the downregulated host proteins in regulation of the cell cycle. The gene annotation analysis was performed on Biological Process and KEGG-pathways terms using the Cluego plug-in on the Cytoscape platform for all genes significantly regulated at 10h p.i. and ordered by decreasing corresponding -log_10_(FDR). (D) Validation of the proteomics analysis set up and results. Representative western blot against viral proteins and two host proteins upregulated in the proteomics at 10h p.i. on the inputs (Total input) and pull- down fractions of AHA-labelled and lysed RAW264.7 cells at 6 and 10h p.i. followed by click reaction and streptavidin pull-down, using AHA-unlabelled cells as a negative control. (E) Comparison of the proteomic and translatomic results in RAW264.7 showing both differential expression and differential protein stability in MNV-infected cells at 10h p.i., with a decrease of stability for all the proteins identified as downregulated in the proteomic analysis. Heatmap of comparison of the log_2_(FC) of the genes identified by proteomic in MNV vs MNV(UVi) (translated proteins) against their corresponding log_2_(FC) in total (Total RNA) and polysomal fraction (Polysomal RNA) generated from the translatomic data. Stars represent significant differential expression as described in Fig. 1A. (F) Validation by western blot of some of the upregulated genes in MNV-infected RAW264.7 cells identified by RNA sequencing but absent from the proteomic dataset. Mock and MNV-infected cells lysates were run on SDS-Page alongside poly I:C treated cells at 20µg/mL and amino acid starved cells for 10h as positive control for the upregulation of TNF and ATF3 respectively.

### Secreted Gdf15 contributes to immunosuppression during MNV infection

Next, we addressed the propagation of an anti-inflammatory signalling from MNV-infected cells to bystander cells after the first round of viral replication. In response to metabolic stress, CHOP and ATF3 upregulate the expression of Gdf15, driving the M2 anti-inflammatory phenotype in macrophages (35). Given that the *Gdf15* transcript is upregulated in MNV-infected RAW264.7 cells as shown on Fig. 3B and C, we performed immunoneutralization assays to evaluate its contribution to MNV propagation. RAW264.7 cells were infected with MNV at different MOI for 24 hours, equivalent to 2 rounds of infection (12h p.i.) and treated with 0.125µg/mL to 1 µg/mL of Gdf15-neutralising antibody before assessing cell survival. The results showed a significant increase in cell survival at the higher dose of the neutralising antibody compared to control IgG, suggesting a correlation between MNV propagation and Gdf15 activity (Fig. 7A). To further characterise the impact of Gdf15 neutralisation, cytokines levels were measured by qRT-PCR in cell cultures inoculated at MOI of 0.1 and 1 at 12h p.i. and after the death of the first round of MNV-infected cells (Fig.7B to E). Gdf15 immunoneutralization resulted in increased levels of the pro-inflammatory *TNF* and *Il-6* transcripts, as well as the STAT5 target genes *Csf2* and *Ptgs2* and potentiation of the paracrine type I IFN response such as the ISG *Mx1* transcript, previously identified as a norovirus restriction factor (54). Overall this suggests that the secretion of Gdf15, downstream of ATF3/CHOP activation, results in immunosuppression and impaired antiviral paracrine signalling to promote MNV propagation.

**Fig. 7:**
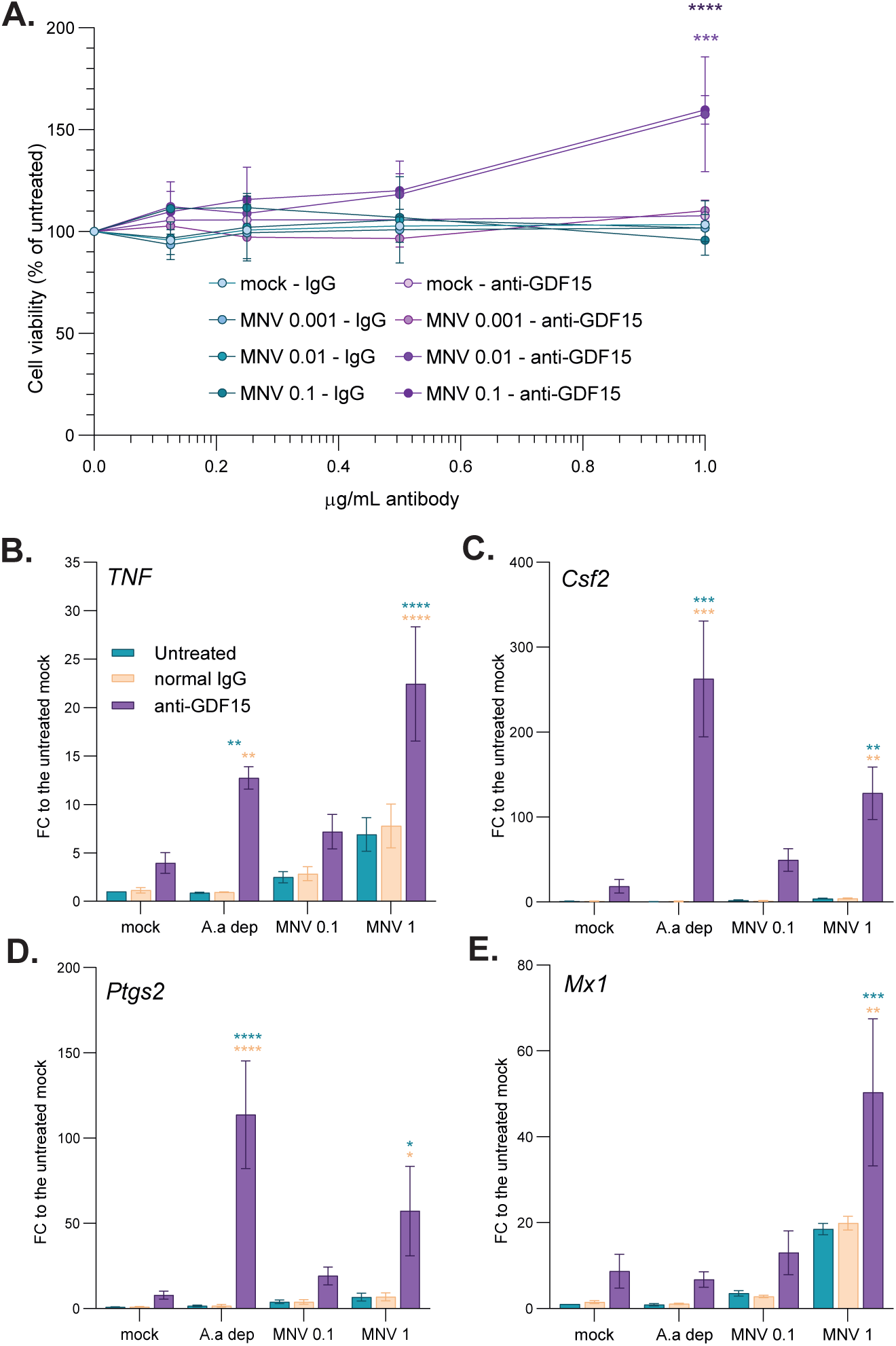
MNV infection induces the secretion of immune-tolerogenic factors. Neutralisation of Gdf15 in the cell culture supernatant resulted in increased cell survival (A) and upregulation of expression of inflammatory and anti-viral factors (B- D). (A) Representative plot (n=4) showing the mean results of cell viability assays performed at 30h p.i. on RAW264.7 cells, mock- or MNV-infected at MOI 0.01, 0.1 and 1 and incubated with normal goat IgG or anti-Gdf15 antibody from 6h p.i. onward at the indicated concentrations. Results are shown as mean ± s.e.m. and statistical analysis results shown above the dots, ** *P*<0.0001, *** *P*<0.001. (B-E) Representative bar plots (n=3) of qRT-PCR analysis of innate immune response transcripts levels. Experiment performed on total transcripts from RAW264.7 cells mock- or MNV-infected at MOI 1 for 12h p.i. and incubated with normal goat IgG or anti-Gdf15 antibody from 6h p.i. onward. Purified RNAs were reverse transcribed using polydT primer and qPCR performed using exon-junction spanning pair of PCR primer. Values are mean ± s.e.m and statistical significance are shown above the bars, **** *P* < 0.0001, *** *P* < 0.001, ** *P* <0.01, * *P* < 0.1. (B) *Tnf* mRNA, (C) *Csf2* mRNA, (D) *Ptgs2* mRNA, (E) *Mx1* mRNA.

## Discussion

Previous genome-wide analyses of the transcriptome reprogramming induced during MNV replication detected perturbations in the innate immune response (40) and differences between cellular models in terms of kinetics of the host response (41). However, both studies are based on comparative analysis of uninfected cells with uninfected controls at 0h p.i., introducing artificial sources of variation and statistical noise. Levenson *et al.* also concluded that upregulation of *IFNβ1* transcript is uncoupled from its protein expression in RAW264.7 cells, which, together with our results on the MNV-induced translational shutoff, suggested a regulation of the antiviral response at the translation stage (19, 22, 23). Therefore, our previous identification of the activation of the eIF2α kinase GCN2 led us to re-address the global early and late host response to MNV replication using a multi-omics approach, including matched non-replicative MNV(UVi) as direct parallel negative control, at each time points. The murine macrophage line RAW264.7 was infected at high MOI (10) and analysed at 6 and 10h p.i., equivalent to the first round of viral replication, to minimise the impact of paracrine immune signalling or apoptotic signalling after the death of the first infected cells (*circa* 12h p.i.). Translational control analysis comparing changes in polysomal against total cytoplasmic fractions at 10h p.i. showed a very high correlation of regulation in both fractions, which fits the known control of expression of the genes associated with the innate immune response. This result seemed to rule out any viral strategy of control at the translational level (12, 55, 56). It also demonstrated proper recruitment of the translational machinery by newly synthesised transcripts thus confirming the absence of effect of P-eIF2α on translation efficiency during MNV replication (23).

More intriguingly, we could not identify an appropriate pro-inflammatory NF-κB response that would normally be instrumental in the secretion of pro-inflammatory autocrine and paracrine cytokines, potentiation of the IFN response and induction of the anti-viral response via activation of JAK and Stat signalling (8, 57–59). This absence of induction of an entire arm of MDA5 response is a crucial aspect of the cellular response to MNV infection, likely to be linked to its ability to replicate and propagate *in vitro* and *in vivo* (20). Remarkably, this seems coupled to the upregulation of stress-related genes such as *ATF4* and *ATF3* and their respective targets *CHOP, Chac1* and *Gdf15*. We propose these characteristics of efficient ISR signalling, downstream of GCN2 activation, and concomitant stimulation of the ATF2/c-jun in response to metabolic stress, could be the key in antagonising the anti-viral inflammatory response (29, 30).

While the intrinsic apoptosis induced by MNV has been shown to favour viral replication by preventing inflammasome related cell death (44), ATF3 is the main driver of the M2 polarisation of macrophages, promoting resolution of inflammation, establishment of adaptive responses to stress and survival (37, 38). ATF3 acts as a secondary modulator of TLR-induced inflammation *in vitro* and *in vivo* via direct transcriptional repression of NF-κB and its target genes (33, 60–62). ATF3 upregulation in response to stress, prior to LPS induction of TLR responses was shown to suppress the upregulation of *TNF* and *Il*-6 (63). This cross-tolerance is part of the immune system homeostasis to avoid excessive and prolonged inflammation. Finally, it is also responsible for the effects of non-steroidal anti-inflammatory drugs via the upregulation of Gdf15 (35).

Time-course analysis of the activation of *ATF3* in MNV-infected RAW264.7 cells showed an upregulation concomitant to the MDA5 response, not secondary to it, which could therefore be responsible for the suppression of NF-κB response downstream of MDA5 activation. Furthermore, we showed by immunoneutralization that Gdf15, direct target gene of both ATF3 and CHOP, is responsible for a paracrine anti-inflammatory signalling in MNV-infected cell cultures. It thus appears that the activation of such a cellular response, stress homeostasis and apoptosis, may have positive effect on replication and viral fitness when triggered at the right time, before the recognition of the viral products by the cell. Too early and the apoptosis will prevent the replication and propagation, too late and the paracrine inflammatory signalling will block it.

The previously identified GCN2 activation along with *Asn* and *Slc1a4* gene upregulation points toward a response to amino acid starvation, and metabolic imbalance in MNV-infected cells. Our characterisation of the level of intracellular free amino acids by LC-MS showed evidence of glutamine depletion as seen by its increased level compared to the mock. Glutamine is one of the critical amino acids involved in central carbon metabolism, forms a large proportion of proteins and participates in immune cell function. Aspartate and glutamate are produced from the TCA cycle metabolism and are the metabolic precursors for the biosynthesis of other proteinogenic amino acids. Increased pool sizes of these amino acids therefore suggested an increased metabolic activity through the TCA cycle, probably required to sustain the cellular metabolic stress and matching the MNV-induced increase of glycolysis previously described (64). It is a possibility that the unchallenged production of viral proteins, depleting the cells in both amino acids and available energy is the main factor responsible for this event. It is also noteworthy to mention that the protein VP2 contains stretches of polyglutamine which are conserved among different strains of MNV, where glutamine on its own represents 12% of amino acid usage against 3.7% in average usage in vertebrates (Fig S5A and B). Interestingly, these polyglutamine motifs also exist in other members of the *Caliciviridae* family (Fig S5C). It is therefore conceivable that the high usage of those amino acids creates elongation disruption leading to ribosome collisions and responsible for both the partial translational shutoff and activation of GCN2 in MNV-infected cells (28).

Overall, we describe the occurrence of a metabolic stress resulting from MNV replication, potentially responsible for GCN2 and ATF2 pathways activation, and the homeostatic response of the cells as driver of the host tolerance to MNV replication and propagation. The hijack of these cellular mechanisms of cross-tolerance and intrinsic apoptosis by MNV by early induction of a metabolic stress seems to be a passive event. But it may also be linked to macrophages in particular and explain the preferential tropism of MNV for cells of the myeloid lineage, in addition to the presence of the CD300 MNV receptor. Finally, this strongly prompts to re-evaluate the role of metabolic stress during viral infection and its impact on the cellular antiviral response. It may also offer an exciting means of increasing the infection and replication of human norovirus *in vitro* by infecting cells subsequent to a transient metabolic stress to induce a cellular tolerance to infection, by incubation in amino acid restricted medium.

## Material and Methods

### Cell lines and viruses

Cells and viruses used in this study were previously described in (23). Briefly, murine macrophage cells RAW264.7 and murine microglial cells BV2 were maintained in Dulbecco’s modified Eagle’s medium (DMEM), 4,5g/L D-glucose, NaPyruvate, L- Glutamine supplemented with 10% foetal calf serum (FCS), 100U of penicillin/ml, 100μg of streptomycin/ml and 10mM HEPES (all supplements purchased from Invitrogen) at 37°C in a 5% CO_2_ environment. Murine norovirus 1 (MNV-1) strain CW1 was described previously (65) and was propagated in BV2 cells as described in with an extra step of concentration using Amicon Centrifugal Filters 100k (Millipore). UV-inactivation was carried by irradiation of virus-containing supernatant in a 15cm plate prior the concentration step in Stratalinker 1800 (Stratagene) at the standard power for 3 times 3min with swirling of the plate between each round. Virus titres were estimated by determination of the TCID_50_ in units per millilitre in RAW264.7 cells and infections were carried at a MOI of 10 unless stated otherwise. The times post-infection refers to the time elapsed following medium replacement after a 1h inoculation period at 37°C in a 5% CO_2_ environment. TLR3 antiviral response was elicited by addition of the synthetic double-strand RNA poly I:C (reconstituted at 10mg/mL in PBS by incubation at 50°C, Sigma) at 20µg/mL directly into the cells medium for the indicated times. For starvation experiment, cells were washed twice in Glutamine, Cysteine and Methionine depleted medium (Invitrogen, thereafter called GCMdep medium) 4.5g/L D-Glucose, supplemented with 10% FBS, 10mM Hepes, 1mM NaPyruvate before incubation for the indicated times in the same medium. GCN2 inhibition was performed by adding A-92 (Axon Medchem) at 500nM to the cells medium for the indicated times.

### Polysomes fractionation and RNA sequencing

Polysomes fractionation has been performed as described in (21). Briefly, 10^7^ RAW264.7 cells uninfected or infected with UVi(MNV) or MNV at MOI 10 for 6 and 10h were treated with 100µg/mL of cycloheximide (CHX-Sigma) for 3min at 37°C to lock the translating ribosomes onto the transcripts before being placed on ice, washed twice with cold PBS(-) (Invitrogen) containing 100µg/ml CHX and lysed on plate with 1mL of cold lysis buffer (15mM Tris-HCl, pH 7.5, 300mM NaCl, 15mM MgCl_2_, 100µg/mL CHX, 1% (v/v) Triton X-100, 200u RNasin (Promega) in RNAse free water (Qiagen)). Lysis was carried on for 5min, the lysates transferred to microcentrifuge tubes and spun at 10 krpm, 3min at 4°C to remove nuclei. The supernatants were aliquoted in 2 tubes for Total and Polysomal RNA isolation. 500µl of each lysate were loaded onto 10mL 10-60% discontinuous sucrose gradients (15mM Tris, pH 7.5, 300 mM NaCl, 15mM MgCl_2_, 100µg/mL CHX, 1mg/mL Heparin) and spun at 38 krpm for 2h at 4°C using the SW41Ti rotor (Beckmann). Gradients were fractionated using the fraction collector FoxyR1 (Teledyne ISCO, Lincoln NE) in 10 fractions of 1mL in 15mL tubes and UV absorbance was monitored at 254nm, recorded using the PeakTrail software and plotted using Graphpad. Total lysates and gradient fractions were denatured in 1.5mL of 7M GnHCl, mixed vigorously and precipitated with 2mL of 100% EtOH at -80°C overnight. RNAs were pelleted at 4,000 rpm for 50min at 4°C before being resuspended in 400µL of RNAse free water, transferred to microcentrifuge tubes and reprecipitated with 40µL of 3M NaOAc pH 5.2 and 1mL of 100% EtOH at -80°C overnight. Samples were centrifuged at top speed in a microfuge for 30min at 4°C, washed once with 75%EtOH, air dried and resuspended in 30µL of RNAse free water before quantification. The fractions containing the polyribosomes were identified by their composition in 40 and 60S ribosomal RNA on 2% agarose gel, pulled together and purified with LiCl precipitation by addition of an equal volume of 7.5M LiCl solution (Ambion), incubation overnight at -20°C, spun at top speed for 30min at 4°C, washed in 75% EtOH, resuspended in 50µL before a second round of LiCl precipitation and eventually resuspended into 30µL of RNAse free water. Three independent replicates of the experiment were performed before control of the purified RNA quality on Bioanalyser (Agilent) as per the manufacturer’s instructions and quantification on Nanodrop. 1µg of RNA per sample were sent to the Oxford Genomics Centre (Welcome Trust Centre for Human Genetics, Oxford, UK) which generated the cDNA libraries with enrichment in polyA-tailed messenger RNA and performed the RNA sequencing on Illumina HiSeq4000 machine, 75bp PairEnds protocol with multiplexing of the samples.

### Analysis of RNA sequencing data

RNA sequencing reads were quality checked via FastQC (v 0.11.4). Reads were mapped to the mouse genome (GRCm38 reference genome, Hisat2 pre-built index files downloaded from ftp://ftp.ccb.jhu.edu/pub/infphilo/hisat2/data/grcm38_tran.tar.gz) using Hisat2 [v 2.0.5] and to the Murine Norovirus genome MNV1-CW1 (DQ285629) coding sequences (ORF1: ABB90153.1, ORF2: ABB90154.1, ORF3: ABB90155.1) using Bowtie2 [v 2.2.5] (68). The function ‘featureCounts’ (69) from the R Bioconductor package ‘Rsubread’ [v 1.16.1] was used to assign mapped sequencing reads to genome features. Genomic features of the host were defined by the tool’s in-built gene annotations for the mouse genome (NCBI RefSeq gene annotations Build 38.1). Host genomic features were annotated using the R package ‘org.Mm.eg.db’ [v 3.7.0], Ensembl (accessed via the R package ‘biomaRt’ [v 2.38.0]) and GenBank (accessed via the R package ‘Annotate’ [v 1.52.1]. The feature count matrix consisted of the set of Host and MNV feature count and genes annotated as rRNA, miRNA, snoRNA, snoRNA or scaRNA were filtered out of the feature count matrix. Filtering of lowly expressed genes was performed after library size normalisation by keeping genes with at least 0.25 CPM in at least 25% of the samples in both Total and Polysomal fractions. Correlation and PCA analysis had been made on the resultant log_2_(CPM) matrix. The degree of correlation between replicates on the Hisat2 / Bowtie was analysed on respective complete feature count matrix using the R function ‘cor’ with the Pearson method and complete linkage hierarchical clustering using the function embedded within the plot generating R CRAN package ‘superheat’ (70). PCA analyses were completed using either the R function ‘pca’ (71) or the R CRAN package ‘pca3d’ (72). Differential expression analysis was performed using the EdgeR quasi-likelihood pipeline (73, 74). Briefly, after scaling normalisation by library size and compositional normalisation using the trimmed mean of M-values (TMM), count data were fitted to a quasi-likelihood (QL) negative binomial generalised log-linear model. Differential expression in selected contrasts was identified using gene-wise Empirical Bayes quasi-likehood F-tests. Benjamini- Hochberg (BH) procedure was used for multiplicity correction of p-values and a threshold of significance at BH p-value <0.1 was selected (75). A final filtering criterion was applied to each individual significant gene using the open-source genome visualisation tool Integrative Genome Viewer IGV hosted by the Broad Institute (https://software.broadinstitute.org/software/igv/home, (76)) and genes displaying poor or aberrant coverage were discarded. Heatmaps of the significant differentially expressed genes were generated on centred and scaled log_2_(CPM) for each gene using the ‘cor’ function with Pearson method and represented with the ‘superheat’ R package after hierarchical clustering. Translational control analysis was completed using the Bioconductor package Anota2seq (39) with default parameters. Genome ontology (GO) enrichment was performed on the web-based software Metascape (https://metascape.org/gp/index.html#/main/step1, (77)) and on the open-source software platform Cytoscape [v 3.8.0] (42) using the plug-in Cluego (78) with the following query settings: GO_Biological Process and KEGG_pathway with pval<10^-5^ only and a *fair* Kappa score of connectivity. The statistical significance of the enrichment was measured using the hypergeometric test implemented with the Bonferroni step down correction of the p-values and grouping of the GO Term based on the highest significance (Kappa score). The corresponding annotations network was generated using a prefuse force directed layout setting. All other charts were generated on Graphpad [v 8.2.1]. Transcription network enrichment was analysed using the WEB-based Gene SeT Analysis Toolkit (http://www.webgestalt.org/, (79)) using the over-representation analysis method and the transcription factor target database MSigDB (80) [v 6.2]. Comparison between studies or cellular backgrounds results were made on the web-based integrative tool for comparing lists JVenn (http://jvenn.toulouse.inra.fr/app/index.html, (81)).

### Phosphoantibody Array

Assays were conducted as described in (21). Briefly, 300µg of protein from mock- and MNV-infected RAW264.7 cells lysed at 2 and 12h p.i. were analysed using the Proteome Profiler Huam Phospho-MAPK Array (R&D Systems), the signal detected on radiographic film (Fuji RX) and quantified using Image J software.

### RNA purification and qRT-PCR

Approximately 2.10^6^ RAW264.7 cells were plated onto 35mm dishes and infected the next day with MNV at a MOI of 10 as described above. Cells were washed once with cold PBS, lysed on plate using 500µl of lysis buffer from the Quick-RNA Miniprep kit (Zymo Research) and RNA subsequently isolated using the same kit according to the manufacturer’s instructions before quantification on Nanodrop. Reverse transcriptions were carried out on 0.5 to 1µg of purified RNA using the Precision nanoScript2 Reverse Transcription kit (Primer Design) and qPCR were performed on 5µL of the cDNA libraries diluted at 1:10 using the Precision Plus 2x qPCR Mastermix (Primer Design) according to the manufacturer’s instructions and the Quant Studio 7 Flex (Applied Biosystems). For validation of the translatomic results, RT-qPCR were carried out on the RNA sequencing samples as mentioned above and results plotted against each other in Graph Pad. Correlation coefficient was calculated on grouped Polysomal and Total data using the Pearson method in GraphPad. Primers are listed in the supplementary data.

### Biorthogonal labelling for characterisation of *de novo* proteome

Stable isotope labelling of amino acids in RAW264.7 cells culture was performed as described in (82) and carried out in high-glucose DMEM lacking arginine and lysine (Sigma-Aldrich) supplemented with dialysed 10% FBS, 1% L-Glutamine, 1x NEAA, 10mM HEPES and 1x penicillin/streptomycin. RAW264.7 cells were maintained in SILAC media supplemented with Light (R0K0), Medium (R6K4) or Heavy (R10K8) Arginine and Lysine (Cambridge Isotope Laboratories) for 5 passages, ensuring complete incorporation of the different isotopes. For labelling of the neotranslatome, 10^7^ SILAC-labelled cells were either mock-, MNV(UVi) or MNV-infected at MOI=10 TCID_50_/cell before replacing with SILAC media without methionine for 30 minutes at 37 °C at 5.5 h and 9.5 h p.i., then labelled with SILAC media containing 1mM L-azidohomoalanine (AHA, Dundee Cell Products) (53, 83) and incubated for 30 minutes at 37°C prior to lysis with 2% SDS in PBS containing 5 U/μl benzonase (Sigma). Cell lysates were heated for 10 minutes at 95°C and centrifuged at 21000 x g for 10 minutes at 4 °C. The supernatant was collected as total cell lysates and 10% of each sample was kept aside as total cell input. After concentration normalisation using BCA analysis (Pierce) following manufacturer instructions, 1.5 mg total cell lysates were diluted to 0.2% SDS, 0.2% Tx100 in PBS, pH 7.4/PI-E, depleted of endogenous biotinylated proteins by incubation with 30 μl streptavidin beads (Pierce 88817) overnight at 4°C before click reaction with 100 μM of biotin-alkyne, 500 μM THPTA (Dundee Cell Products), 100 μM CuSO_4_, 5 mM Aminoguanidine and 2.5 mM Sodium Ascorbate the reaction allowed to proceed for 6h at room temperature before removal of the biotin excess using Zeba desalting columns (7kDa cut-off, Thermofisher). The samples were then combined altogether at a 1:1:1 ratio within one replicate. Three independent replicates were made with a different SILAC medium assigned to a given condition for each replicate. AHA-labelled biotinylated peptides were affinity enriched by incubation with 300 μl streptavidin beads (Pierce) overnight at 4°C with rotation in PBS containing 0.1%SDS (v/v) and washed 20 times with 0.5 ml cold PBS 1%SDS with complete removal of the liquid before elution of the bound fractions in 75 μl PBS 2%SDS, 1mM Biotin and denaturation at 95°C for 5min. Validation IP was performed similarly but the AHA-labelling time was increased to 2 hours to improve labelling and enrichment. Eluted fractions were submitted for mass spectrometry analysis at the University of Bristol Proteomics Facility as described in (82). SILAC pooled samples were run on an SDS-PAGE gel and each gel lane cut into 10 equal slices. Each slice was subjected to in-gel tryptic digestion using a DigestPro automated digestion unit (Intavis Ltd.) to minimise manual handling. The resulting peptides were fractionated using an Ultimate 3000 nano-LC system in line with an Orbitrap Fusion Tribrid mass spectrometer (Thermo Scientific). In brief, peptides in 1% (vol/vol) formic acid were injected onto an Acclaim PepMap C18 nano-trap column (Thermo Scientific). After washing with 0.5% (vol/vol) acetonitrile 0.1% (vol/vol) formic acid peptides were resolved on a 250 mm × 75 μm Acclaim PepMap C18 reverse phase analytical column (Thermo Scientific) over a 150 min organic gradient, using 7 gradient segments (1-6% solvent B over 1min., 6-15% B over 58min., 15-32%B over 58min., 32-40%B over 5min., 40-90%B over 1min., held at 90%B for 6min and then reduced to 1%B over 1min.) with a flow rate of 300 nl min^−1^. Solvent A was 0.1% formic acid and Solvent B was aqueous 80% acetonitrile in 0.1% formic acid. Peptides were ionized by nano-electrospray ionization at 2.0 kV using a stainless steel emitter with an internal diameter of 30 μm (Thermo Scientific) and a capillary temperature of 275°C.

All spectra were acquired using an Orbitrap Fusion Tribrid mass spectrometer controlled by Xcalibur 2.1 software (Thermo Scientific) and operated in data- dependent acquisition mode. FTMS1 spectra were collected at a resolution of 120 000 over a scan range (m/z) of 350-1550, with an automatic gain control (AGC) target of 3E5 and a max injection time of 100ms. Precursors were filtered using an Intensity Range of 1E4 to 1E20 and according to charge state (to include charge states 2-6) and with monoisotopic precursor selection. Previously interrogated precursors were excluded using a dynamic window (40s +/-10ppm). The MS2 precursors were isolated with a quadrupole mass filter set to a width of 1.4m/z. ITMS2 spectra were collected with an AGC target of 2E4, max injection time of 40ms and CID collision energy of 35%.

### Analysis of SILAC – AHA proteomics data

The raw data files were processed and quantified using Maxquant software v1.6.1.0 and searched against the Uniprot Mouse database (downloaded 18/03/30: 61314 entries) and MNV1 protein sequences (DQ285629.1). Peptide precursor mass tolerance was set at 20ppm, and MS/MS tolerance was set at 0.5Da. Search criteria included oxidation of methionine (+15.9949), acetylation of protein N-termini (+42.0106), Triazole-PEG4-biotin of methionine (+452.238Da), AHA of methionine (- 4.9863) and SILAC labels (+6.020Da and +10.008Da at arginine and +4.025Da and +8.014Da at lysine) as variable modifications. Searches were performed with full tryptic digestion and a maximum of 2 missed cleavage was allowed. The reverse database search option was enabled and the data was filtered to satisfy false discovery rate (FDR) of 1%.

Differential expression was analysed by computing the pairwise ratios of either pair of mock-, UVi(MNV) and MNV- infected samples, the Log_2_SILAC ratios for proteins identified in at least 2 out of 3 replicates were averaged and proteins considered significantly regulated if the pairwise ratio was in the top or bottom 2.5% either contrasts MNV:mock or MNV:UVi(MNV). Subsequent computational analysis had been made as described in the above section “Analysis of RNA sequencing data”. The results were validated against the total inputs by immunoblotting for some of the major proteins identified as significantly enriched in the *de novo* proteome at 6 and 10h p.i.

### Immunoblotting

Immunoblotting analysis was performed as described in (23). Briefly, approximately 2.10^6^ RAW264.7 cells were plated onto 35mm dishes and infected the next day with MNV at a MOI of 10 as described above. At the indicated times, cells were lysed in 150μL of 1x Gel Loading Buffer (New England Biolabs), sonicated and boiled 5min at 95°C. Cell lysates were separated by SDS-PAGE, using 10μg of total proteins, and the proteins transferred to polyvinylidene difluoride membranes. These were then probed with the following primary antibodies: mouse anti-Phospho-ATF2 T71 (1:1,000, Abcam), rabbit anti-ATF3 (1:500, Cell Signalling), goat anti-TNFα (1:500, Santa Cruz), rabbit anti-Ddx58 (1:500, Santa Cruz sc-376845), rabbit anti-Stat1 (1:1000, Abcam Ab92506), mouse anti-VP1 (1:500, Prof. Goodfellow, Cambridge, UK), anti-MNV-3 mouse immune sera (1:1,000 Prof. Goodfellow, Cambridge, UK) rabbit anti-NS7 (1:10,000, Prof. Goodfellow, Cambridge, UK), mouse anti-GAPDH (Clone 6C5, 1:20,000, Invitrogen); followed by incubation with the appropriate peroxidase labelled secondary antibodies (Dako) and chemiluminescence development using the Clarity Western ECL Substrate (Bio-Rad). The results were acquired using the VILBER imaging system.

### Quantification of total amino acids

2.10^6^ RAW264.7 cells were plated on 35mm plates and Mock or MNV-infected the next day at MOI 10 or washed once and incubated in GCM-dep medium instead of the normal medium for the cells in starvation control. At 10h p.i., cells were placed on ice, scraped in cold PBS, transferred to a 1.5mL tube, washed one more time in cold PBS and the pellets kept at -80°C. The pellets were then resuspended in 300µl of cold L-Amino Acid Assay buffer according to the manufacturer instructions (L-Amino Acid Quantification kit, Sigma) and lysates were then centrifuged at 13,000 rpm for 10min at 4°C to remove insoluble material. The supernatants were transferred to a new tube and the amino acids concentrations measured in duplicate on 50µL of each lysate by adding one volume of the Master Reaction Mix and 30min incubation at 37°C in the dark. Results were measured at 570nm in ClarioStar plate reader against the kit control standard curve and normalised by the protein contents quantified using the BCA kit (Pierce) according to the manufacturer’s instructions.

### Quantification of free individual amino acids

6.10^6^ RAW264.7 cells per condition were plated onto 15cm plates and treated as mentioned above. At 10h p.i., metabolites were harvested by collecting the cells in cold PBS, followed by two washes in cold PBS, resuspension in 400µL of TritonX100 0.1% (v/v) in LC-MS grade water (Fisher) and lysis in 2:1 methanol (400µL): chloroform (200µL) (LC-MS grade Fisher). Lysates were mixed well and incubated 10min on ice. The lysates were then extracted by incubation 10min at room temperature (RT) before phase separation by centrifugation at 8krpm for 10min at RT. Metabolites were separated into polar and non-polar phases. The free amino acids were recovered in the polar phase and used for the metabolomics analysis by Gas-Chromatography mass spectrometry (GC-MS). The intermediate phases containing the proteins were used for determination of the protein content after being spin down at full speed for 10min, complete removal of the liquid phases, air-dried for 5min before lysis in 300µL of 0.1N NaOH and sonication until clear. Quantification of the protein contents was made using the BCA kit according to the manufacturer’s instruction. Three independent replicates were completed before being further processed by Gas-Chromatography mass spectrometry (GC-MS). Authentic standards for 21 amino acids were purchased from Sigma Aldrich and were used for method development. Norvaline (Sigma Aldrich) was used as the internal standard for GC-MS analysis and for data normalization. 5µL of 10 ng.µL^-1^ norvaline was added to the extracted polar phase and dried. Standards and dried metabolite extracts were mixed with 25 µL pyridine and incubated at 900rpm, 37°C and for 30 minutes. 35 µL tert-butyldimethyl silyl chloride (TBDMSCl) (Sigma Aldrich) was added to the pyridine-metabolite extracts and were mixed at 900rpm, 60°C and for 30 minutes. After this incubation, samples were briefly spun, transferred to GC-MS vials and sealed. 1 µL of the derivatized metabolites were injected into an injection port of GC 7890 system temperature maintained at 230 °C. Amino acids were separated using a VF-5ms inert 5% phenyl-methyl column (Agilent Technologies). The oven temperature was constant at 120 °C for 5 min, followed by an increased to 270 °C at 4 °C/minute and was held at this temperature for 3 min. The temperature was further increased to 310°C at 20 °C/minute and was held at this temperature for 1 minute. The carrier gas was helium and the flow was maintained at 1.3 mL/minute. The total run time of each sample was 51.667 min, including a solvent delay of 10 min. The scan parameters for the MS detector were set to 150 for low mass and 600 for high mass. MS data was detected using electron ionisation (EI) on a 5795 MS system with triple axis detector (Agilent Technologies). The scan parameters for the MS detector were set to 50 for a low mass and 600 for a high mass. Identification of free amino acids was done in AMDIS (Automated mass spectral deconvolution and identification system) developed at the National Institute of Standards and Technology (NIST) and were identified by their characteristic fragmentation patterns and m/z values as previously published (84). GC-MS data was extracted using the Chemstation software (Agilent Technologies) and detection and quantitation parameters previously established (84). Total ion counts for the M-57 m/z fragments from 21 amino acids were extracted for quantitation.

### Cell viability assay

RAW264.7 cells were Mock or MNV-infected at MOI 0.1, 0.01 and 0.001 in complete medium and 5.10^3 cells in 100µL /well were plated onto a white walled and clear bottom 96 wells plate (Costar) in duplicate. Neutralising goat antibody against human GDF15-or normal goat IgG (R&D systems) was added at 6h p.i. at 0.125, 0.25, 0.5 and 1µg/mL and the cells incubated until 24h p.i., ensuring 2 rounds of infection, and the plates kept at -80°C until further processing. Cell viability was assessed using the CellTiter-Glo 2.0 assay (Promega). Briefly, 100µL of room temperature reagent was added to each well on ice and mixed by pipetting. The plates were then incubated at room-temperature for 1h before reading of the luminescence on the BMG microplate reader (Labtech) using the Clariostar software with an integration time of 2s. The results were blank adjusted and normalised against the untreated cells of the same condition.

### Accession number

The raw RNA-seq data and mapping results can be found as part of GEO repository database, submission GSE168402. The raw proteomic data, search results and FASTA file can be found as part of PRIDE repository database, PXD025169.

## Supplementary information captions

**S1 Fig: Polysomes fractionation (A-C) and exploratory analysis of the RNA sequencing data in RAW264.7 cells (D to I).** (A-C) Absorbance profiles of the total cytoplasmic extracts fractioned onto 10 to 50% sucrose gradients, monitored at 254nm and recorded on PeakTrail for the three biological replicates used in this work. MNV(UVi) 6h p.i. (light blue), MNV(UVi) 10h p.i. (cyan), MNV 6h p.i. (light purple), MNV 10h p.i. (deep purple). Fractions 1 to 2, free material; fractions 2 to 4, free ribosomal subunits 40 and 60S and monosomes 80S; fractions 5 to 10, polysomal fractions. Note the decrease in polysomal material in MNV 10h p.i. samples compared to the MNV(UVi) counterpart matching the translation shutoff previously observed in MNV-infected cells after 6h p.i. (23). (D) Bar plot of the total number of reads for each sample (blue bars) and corresponding number of reads mapped to the host genome (GRCm38.p5 murine genome primary assembly) using Hisat2 (green bars) software. (E) Scatter plot showing the percentage of total reads mapped to the murine genome using either Hisat2 (blue dots, left *y* axis) for each sample and the subsequent percentage of reads mapping to MNV1-CW1 genome (DQ285629) (purple dots, right *y* axis). (F) Mapped reads using Hisat2 were assigned a genomic feature (complete dataset) subsequently annotated using GenBank. All genes annotated as rRNA, miRNA, snoRNA, scaRNA were filtered out (filtered dataset). (G) Analysis of the batch effect of the biological replicates on the RNA sequencing outputs measured by calculation of the pair correlation of all samples from the complete Hisat2 (Host) and Bowtie (MNV) dataset, represented as a matrix. Correlation calculation using the Pearson method on log_2_CPM of the complete dataset filtered for low reads genes and small non-coding RNA, coupled with complete linkage hierarchical clustering showing the low impact of the batch effect with high correlation for samples of the same condition across the three replicates (average *r* = 0.979 ± 0.007) and highlighting the multiple sources of variation between the samples as infection, time post-infection and total or polysomal fraction. (H – I) 3D representation of principal component analysis of the complete Hisat2 (host) and Bowtie (MNV) dataset (H) or the Hisat2 only (host) dataset (I) showing the combinatorial order of the variation parameters in presence or absence of the viral RNA in the dataset. Principal components scores were calculated on log_2_CPM datasets filtered for low reads genes and small non-coding RNA. 3D cluster plot of the results showed grouping of the samples following the variables identified by correlation analysis and a clear separation of the samples from MNV-infected cells lysed at 10h p.i. independently of the occurrence of the viral RNA in the dataset, highlighting the combination “infection” and “10h p.i.” as the main sources of variation in this experiment.

**S2 Fig.: Differential expression analysis of MNV-infected cells compared to MNV(UVi)-infected cells in overtime comparison exposed wide array of changes common to both conditions.** Scatter plot of the results of the multivariate analysis of the differential expression comparing the 10 vs 6h p.i. time points for the MNV-infected cells vs the MNV(UVi)-infected cells in the Total fraction (A) and the Polysomal fraction (B). Results were expressed as log_2_ (FC 10vs6) and gene changes were considered significant when associated with a BHpval <0.1. MNV and MNV(UVi) results were merged, omitting genes identified in only one condition, and their respective log_2_(FC) for the significant dataset were used in the biorthogonal projection, MNV-infected cells (*y* axis) vs MNV(UVi)-infected cells (*x* axis). Genes were further subdivided in 3 groups: significant in MNV-infected cells only (green dot), significant in MNV(UVi)-infected cells only (purple dot), significant in both (orange dot). (A) Comparison in the total fraction showing positively correlating regulated genes in both contrasts (linear regression, R^2^=0.9135, pval<0.0001). The apparent trend of genes regulated only in MNV(UVi) clustering in and near the 95% confidence prediction bands of linear regression from the genes regulated in both contrasts suggested a stringent significance threshold rather than an actual difference between the conditions. The genes regulated in MNV-infected cells only can be subdivided into 3 clusters: in and around the IC95%, an important cluster of upregulated genes and a small cluster of downregulated genes, both likely to be MNV-specific. (B) Same comparison than in A. for the RNA from the Polysomal fraction. (C) Heatmap of the comparison of the significant differentially expressed genes in MNV-infected cells overtime (green dots in A. and B.) displaying their log_2_CPM values in each sample showing an absence of strict clustering following infection, time, fraction or replicate suggesting the predominance of uncontrolled variability sources and unsuitability of this analysis design. Values were centred and scaled before hierarchical clustering using Pearson method and complete linkage hierarchical clustering. (D) Scatter plot of the differential expression analysis of the contrasts MNV-infected cells vs MNV(UVi)-infected cells at 6h p.i. in Total (*x* axis) vs Polysomal (*y* axis) fraction showing no significant changes. (E) Validation by qRT-PCR of the translatomic results at 10h p.i. Scatter plot of the log_2_FC from the differential expression analysis (green dots) for the contrasts MNV-infected cells vs MNV(UVi)-infected cells at 10h p.i. in Total (*x* axis) vs Polysomal (*y* axis) fraction compared with the log_2_FC obtained by qRT-PCR (purple dots) showing a high degree of correlation (Spearman ρ= 0.9564, pval<0.0001).

**S3 Fig.: MNV-induced upregulated genes belong to the IFN response and to the cellular response to stress linked to intrinsic apoptotic pathway.** Annotation network of the GO analysis of the genes significantly regulated in MNV-infected cells (Total and Polysomal) at 10h p.i. as presented in Fig. 2F and generated using a prefuse force directed layout setting.

**S4 Fig.: Neotranslatome analysis at 6h p.i.** RAW264.7 cells proteins were subjected to labelling with media supplemented with Light (R0K0), Medium (R6K4) or Heavy (R10K8) Arginine and Lysine for at least 5 passages before assessment of their suitability to the proteomics analysis. Biorthogonal projection of the proteomics analysis results plotting the genes significant (blue dots) in both contrasts for cells mock-, MNV(UVi), or MNV-infected at 6h p.i., linear regression analysis showing a high level of correlation (*Pearson r* =0.9727, pval<0.0001).

**S5 Fig: Murine Norovirus small basic protein VP2 is enriched in glutamine residues.** (A) Analysis of the amino usage in MNV1 CW1 proteins (polyprotein YP_720001.1 purple, VP1 YP_720002.1 pink, VP2 YP_720003.1 blue, VF1 YP_006390081.1 green) showed enrichment of VP2 in glutamine compared to the average usage in vertebrate (grey) and other MNV1 proteins (B). Conservation analysis between murine norovirus strains showing a very high level of conservation of VP2. (C) Protein superfamily analysis using the Conserved Protein Domain Family tool in NCBI on pfam 03035 (RNA_capsid) showing a partial conservation of the glutamine usage in VP2 of other *Caliciviridae* family members.

## Supplementary data

Table 1: Results of the differential expression analysis of the RNAseq experiment performed in RAW264.7 cells at 10h p.i.

Table2: qPCR primers list.

## Acknowledgement

Work in N.L.’s laboratory is supported by Biotechnology and Biological Sciences Research Council research grant number BB/N000943/1. I.G. is a Wellcome Senior Fellow [Ref: 207498/Z/17/Z] and also supported by the Biotechnology and Biological Sciences Research Council [grant number BB/N001176/1].

